# Pancreatic α-cells are required for nutrient homeostasis by regulating dynamic β-cell networks in islets

**DOI:** 10.64898/2026.03.02.709124

**Authors:** Marie Lallouet, Manon Jaffredo, Antoine Pirog, Karen Leal-Fischer, Julien Gaitan, Daniel T. Meier, Sylvie Renaud, Matthieu Raoux, Jochen Lang

**Affiliations:** Univ. Bordeaux, CNRS, Bordeaux INP, CBMN, UMR 5248, Institut de Chimie et Biologie de Membranes et des Nano-objets, CBMN, UMR 5248, F-33600 Pessac, France; Electronics-Physics-Acoustics Department, Junia, Lille, France; Univ. Bordeaux, CNRS, Bordeaux INP, Intégration de Matériaux dans les Systèmes, IMS, UMR 5218, F-33400 Talence, France; University of Basel, Department Biomedicine, Basel, Switzerland

## Abstract

Pancreatic islets contain α-, β-, γ- and δ-cells as sensors and actuators regulating glucose homeostasis. Despite the known importance of α-cells, they are seemingly required for glucose tolerance only under metabolic stress. In an inducible model of α-cell ablation in mice (GluDTR), glucose tolerance was considerably decreased by physiological addition of amino-acids mimicking meals. Analysis of islet β-cell secretion and electrical activities using microelectrode arrays (MEA) detected only minor differences in GluDTR mice for glucose but revealed a major reduction upon addition of amino acids. Analysis of functional islet β-cell networks by high density MEA revealed leading regions in different locations, a high degree of synchrony and the activation of large cell clusters. The characteristics of leading regions were preserved in GluDTR islets, but synchrony, cluster size and signal propagation speed were largely reduced. Thus, even without metabolic stress, α-cells are required for nutrient homeostasis by regulating the dynamics of β-cell networks.

**Teaser:** Islet α-cells are required for meal tolerance by adjusting synchrony, cluster size and signal propagation of β-cell networks.

## Introduction

Pancreatic islets play a major role in nutrient homeostasis and their dysfunction leads to the different forms of diabetes (1–2). These micro-organs consist of four major cell types: the insulin secreting β-cells and proglucagon derived peptides (glucagon, GLP-1) secreting α cells release hormones into the systemic circulation, whereas somatostatin secreted from δ-cells and the less explored γ-cells are thought to have a role mainly within the islets (3). Islet cells have the simultaneous role of actuators and sensors: whereas β-cells mainly sense glucose, α-cells are important for glucose and for amino-acid detection (4–5). The adaptive function of islets requires crosstalk between the different cell types (6) and activities of α and β-cells are known to oscillate in a phase-locked fashion in-vivo and in-vitro (7–9). Islet α-cells release glucagon and a minor amount of GLP-1 (10–13). Whereas glucagon can activate both, glucagon and GLP-1 receptors, GLP-1 is specific and far more potent for its cognate receptor (12). Both receptors activate downstream adenylyl cyclases as well as phospholipases (14–16) and the effects of glucagon as well as GLP-1 on insulin secretion from β-cells are mediated by both the cAMP-dependent and cAMP-independent pathways (17–18).

Simulations of α-β crosstalk predict that a specific model, where α-cells activate β-cells and β-cells inhibit α-cells, results in the most harmonic response to changes in glucose concentrations with improved glucose tolerance and amino acid handling (19–20). Nevertheless, the precise interaction between islet cell types under various nutrient stimulations is still not fully elucidated. Mouse models with ablation of a given cell type via specific expression of the diphtheria toxin receptor (DTR) and treatment with diphtheria toxin have provided information on cell-type specific in-vivo interaction. The deletion of α-cells (GluDTR) did not significantly alter glucose tolerance in young mice and led to an impairment only upon metabolic stress or ageing (21–23) similar to the observations in mice lacking the β-cell glucagon and GLP-1 receptors (10, 24). Moreover, deleting all cell types except for β-cells improved glucose tolerance in-vivo with normal insulin secretion in-vitro (25). In-vitro analysis of GluDTR mice showed a reduced in-vitro insulin secretion in response to glucose that could be restored by addition of exogeneous glucagon, GLP-1 or the adenylyl cyclase activator forskolin (12, 24). Similarly, the effect of amino acids on insulin secretion was diminished in-vitro in a mouse model where the α-cell activity was either blunted by activation of an inhibitory designer G-protein coupled receptor (26) or the glucagon receptor was absent on β-cells (10).

Amino acids account for one third of the calories in a standard meal, they increase insulin secretion in humans and lower blood glucose (27). In recent years, the awareness has increased about the role of α-cells as sensors for amino acids (28) and the influence of amino acids on the secretory activity of β-cells via α-cells in-vitro has been established (26). However, their effect on cellular or network properties have not been addressed. β-cells are electrogenic and their increase in metabolism leads to changes in plasma membrane ion fluxes resulting in insulin exocytosis via Ca^2+^-dependent processes (29). Activation of islet β-cells implies dynamic networks and functional coupling among them for an optimal physiological response (30). It remains largely unknown how these membrane ion fluxes and electrical networks are modulated by α-cells during the long postprandial period. Extracellular electrophysiology offers a convenient mean for on-line, non-invasive, and long-term monitoring and does not require biasing methods such as loading of fluorescent agents or manipulations of the sample for genetically encoded sensors. Moreover, electrophysiology offers a temporal resolution unmatched by other approaches. Microelectrode arrays (MEAs) measure changes in field potentials and intercellular coupling. Coordination between islet β-cells, a hallmark of islet activation, can be reliably detected and analyzed by the monitoring of so-called slow potentials without bias (31–37).

We have now used this approach to characterize the roles of glucose and amino acids in β-cell activation, their network dynamics and the role of α-cells herein using acute α-cell specific deletion by diphtheria toxin receptor expression and toxin treatment (23–24). Moreover, we developed the use of high-density MEA for the fine analysis of β-cell activation in islet subregions and to establish the characteristics of physiological β-cell networks in the presence and absence of α-cells. Our in-vivo characterization and islet analysis demonstrate a major role for α-cells in the β-cell response to amino acids in terms of glucose tolerance, islet activity and β-cell network properties.

## Results

### α-cells mediate the effect of amino acids on glycemia and insulinemia

To obtain mice with islets lacking α-cells, we used a genetic mouse model of inducible α-cell ablation subsequent to cell specific expression of the human diphtheria toxin receptor under the control of a glucagon promoter (23–24), termed GluDTR. Injection of diphtheria toxin leads to progressive loss of α-cells (Fig. 1A). Induced GluDTR mice gained some more weight as compared to WT (Fig. 1 B). Islet hormone analysis confirmed a large reduction in glucagon content in GluDTR mice and values were at the limit of detection whereas insulin contents did not differ (Fig 1C). Intraperitoneal glucose tolerance tests demonstrated no difference between WT and GluDTR mice when injecting glucose only (Fig. 1 D, E). Adding a physiological mix of 19 amino acids to glucose led to a significant reduction of glycemia in wild-type (WT) mice and the areas under the curve during the first 30 min were significantly different (28.6±1.2 vs 23.0±0.7; Tukey 2p <0.01). This glycemia-lowering effect of AAM was absent in GluDTR mice (Fig. 1D). To rule out that this was due to an increase in calories injected, we also tested ip glucose tolerance at the same number of calories as glucose and AAM combined, using 3g/kg of glucose (Fig. 1 F, G). Under this condition, glucose tolerance in WT mice again resembled that of GluDTR mice ruling out calories as the responsible factor. As the presence of AAM considerably improved glucose tolerance in WT mice during the first 30 min, we also measured blood insulin and C-peptide levels (Fig. 1, H-K). After ip injection of glucose alone there was no difference in insulin levels between WT and GluDTR mice. The presence of AAM considerably increased hormone levels in WT animals but had no effect in GluDTR mice. Pyruvate tolerance tests provide some crude means to assess the role of liver in glucose homeostasis (38) and glucagon regulates gluconeogenesis (39). This test did not show any difference between WT and GluDTR mice (Fig. 1L). Moreover, fasting insulin or C-peptide levels or insulin tolerance test did not show any difference (Fig. 1, M and N). Thus, the main difference in-vivo between WT and GluDTR animals resides in the glycemic response to amino acids in the presence of glucose.

**Figure 1:**
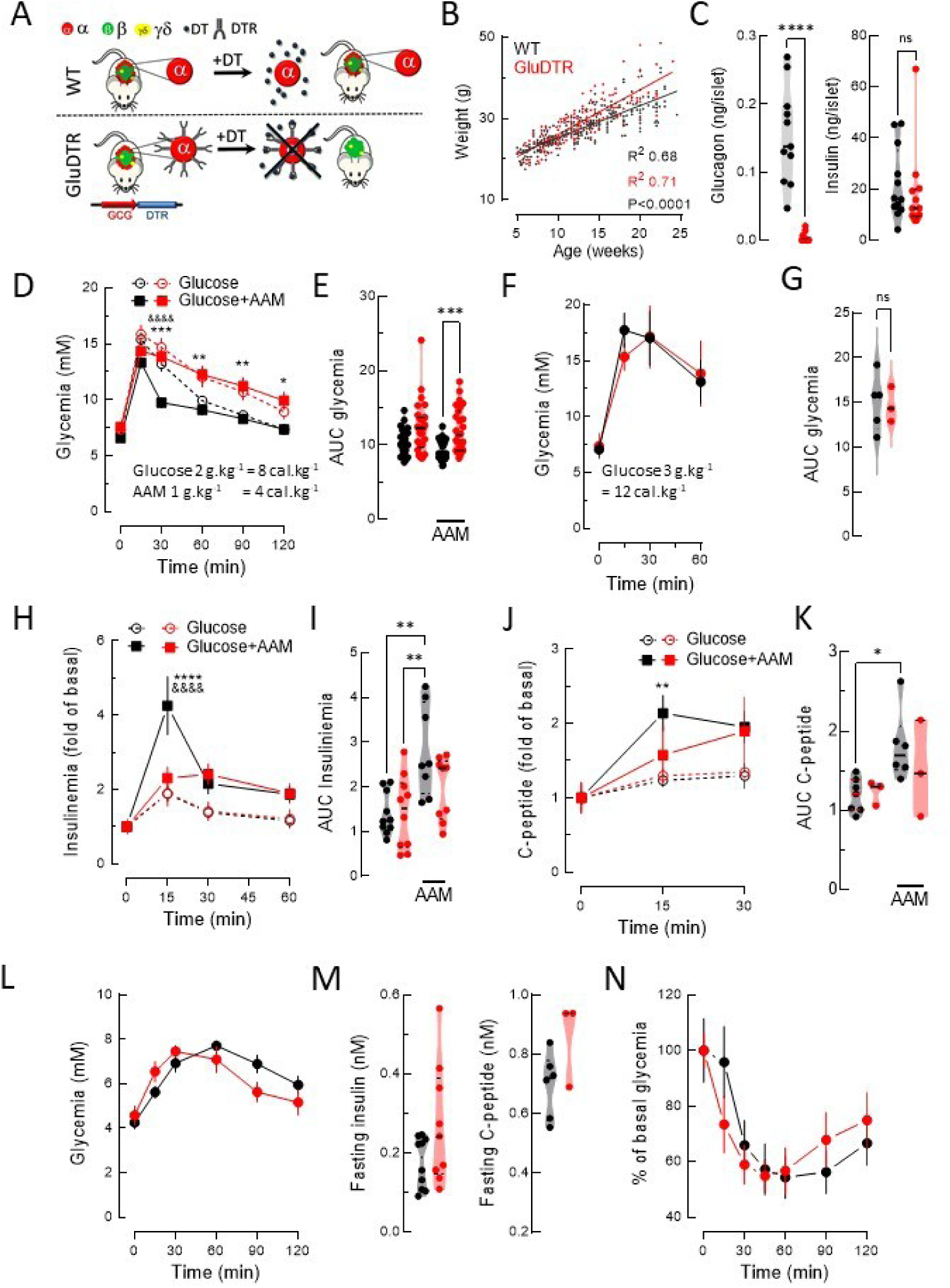
Animal model and in-vivo characterization of WT and GluDTR mice. WT data in black, GluDTR data in red. (**A)** GluDTR mice expressing the human diphtheria toxin (DT) receptor (DTR) under the control of the rat glucagon promoter were fully backcrossed to a C57BL/6N genetic background. Injection of DT at 5 to 8 weeks of age leads to selective destruction of α-cells. (**B)** Growth characteristics in terms of weight. N= 31 – 33. (**C**) Islet hormone contents, N= 10-11. (**D**) Intraperitoneal glucose and amino acid tolerance test in WT and GluDTR mice (2g/kg of glucose and 1kg/g amino acids); AAM, glucose and amino acid mix; N=24-30. € Area under the curve (AUC) of D. (**F**) Intraperitoneal glucose tolerance test in WT and GluDTR mice (2g/kg of glucose). N=3-5. (**G**) AUC of F. (**H and I)** Insulinemia and AUC during intraperitoneal glucose and amino acid tolerance test (see D and E). N=8-10. (**J and K)** C-peptide levels and AUC during intraperitoneal glucose and amino acid tolerance test (see D and E). N=3-7. (**L)** Pyruvate tolerance test (2g/kg pyruvate ip). N=7-8. (**M**) Fasting insulin and C-peptide levels; N=9-10. (**N**) insulin tolerance test (0,5U/kg); N=9-10. *, **, ***, ****, 2p<0.05, 0.01, 0.001 and 0.0001 respectively; in D and H: **** or ***, WT glucose and AAM as compared to WT glucose alone; ^&&&&^, 2p< 0.0001, WT as compared to glucose and AAM in GluDTR. One or 2-way ANOVA, Tukey or Holm-Sidak post-hoc.

### α-cells enhance β-cell electrical activity mainly during the 2^nd^ phase in the presence of AAM

Extracellular electrophysiology, as done here using microelectrode arrays (MEA), provides direct information on the electrical activity of islets and the amplitude of so-called slow potentials (SP) provides an excellent measure of islet β-cell coupling (33, 36). We have used this approach here to evaluate the effect of different glucose concentrations, either alone or in combination with AAM, on islets (Fig. 2). We choose 3 mM glucose (G3) concentrations as basal, 6 mM (G6) as intermediary where islets start to respond and 8.2 mM glucose (G8.2), near to the maximum effective concentration of 10 mM (36). At G3 GluDTR islets were slightly less active in terms of frequency and this was further accentuated in the presence of glucose and AAM. G6 induced a clear 1^st^ and 2^nd^ phase in the frequency and amplitude of SPs (Fig. 2 A, D) without any significant differences between WT and GluDTR islet (Fig. 2C, F).

**Figure 2:**
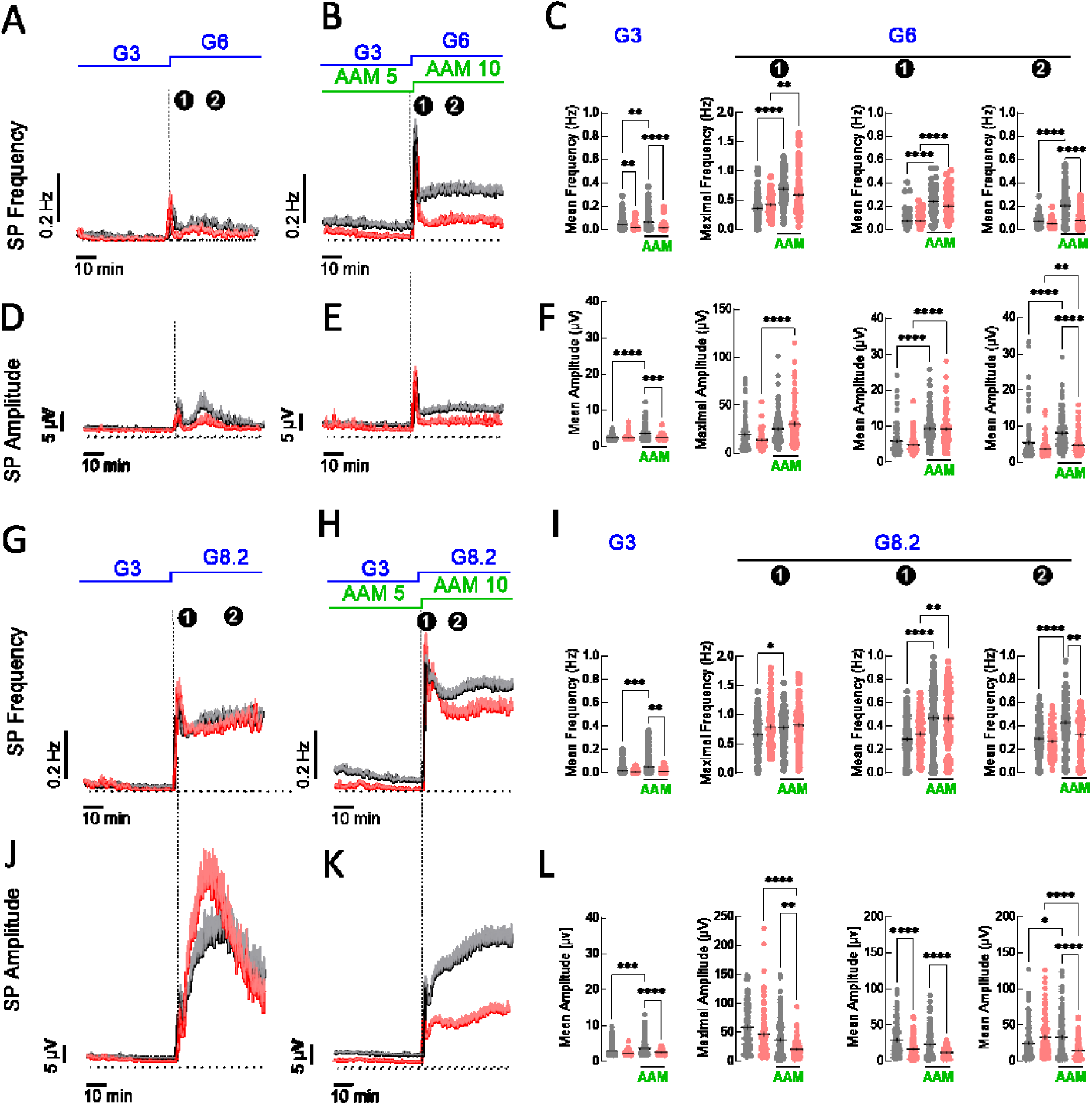
Effects of amino-acids on glucose-evoked electrical activity in islets recorded by MEAs. WT data in black, GluDTR data in red. (**A, B)** Slow potential mean frequencies (±SEM) evoked by 3 followed by 6 mM glucose (G3, G6) in the absence or presence of 5 and 10 mM amino acid mix (AAM5, AAM 10), respectively. 1^st^ and 2^nd^ phases are indicated (❶,❷). N=3-4, n=98-110. **C**: Statistics of A and B (G3; ❶, 0-300 s; ❷, 700-2700 s after increase of glucose) and maximal frequencies of 1^st^ phase. (**D, E, F**) Slow potential amplitudes and statistics as in A, B and C. N, n as in A and B. (**G, H, I)** Same as A to C, mean slow potential frequencies but using 8.2 mM glucose as stimulatory glucose concentration. N=7, n=99-134. (**J, K, L)** Same as D to F, mean slow potential amplitudes but using 8.2 mM glucose as stimulatory glucose concentration. N=7, n=99-134. ANOVA and Kruskal-Wallis, *, **, ***, ****, 2p<0.05, <0.01, <0.001, <0,0001.

In contrast, the addition of AAM to G6 revealed clear differences in WT vs GluDTR islets (Fig. 2 B, E). The presence of AAM increased in both, WT and GluDTR, the mean frequency and amplitude during glucose stimulation in the 1^st^ and 2^nd^ phase. Strikingly the increase in the 2^nd^ phase mean frequency and amplitude was far more pronounced in the WT islets and significantly different from GluDTR islets (Fig. 2 B, C, E, F), the latter arising to only 38 % (frequency) and 59 % (amplitude) of WT islet values. We next examined the effect of a slightly higher glucose concentration, 8.2 mM. In the overall, frequencies and amplitudes were increased as compared to G6. Again, frequencies and amplitudes were comparable between WT and GluDTR islets (Fig. 2 G, J). In the presence of AAM and G8.2 (Fig. 2 H, K) differences were apparent between WT and GluDTR islets in the mean frequencies of the 2^nd^ phase and even more pronounced in mean amplitudes during the 2^nd^ phase (Fig. 2, I, L) which amounted in the GluDTR islets only to 45% of those in the WT. Thus, similar to in-vivo data, the presence of amino acids induced a pronounced difference between the WT vs GluDTR and was present mainly during the second phase.

As the lack of effects of AAM in the absence of α-cells are most likely due to the absence of α-cell hormones, we tested next whether exogenous glucagon can restore the activity of GluDTR islets. To this end we added glucagon to the 2^nd^ phase during stimulation of islets by G8.2 and AAM (Fig. 3). We again observed a marked difference between WT and GluDTR islets upon exposure to glucose and amino acids, especially during the 2^nd^ phase (Fig. 3, A, C). The addition of exogeneous glucagon (Fig. 3, A-D) induced a recovery by increasing the 2^nd^ phase frequencies from 45±5% of WT (both without glucagon) to 92±4 % of WT (both with glucagon). Amplitudes were recovered to a slightly lesser degree from 48±6 % of WT (without glucagon) to 80±7 % of WT (both with glucagon). In contrast, forskolin, an activator of adenylyl cyclases, increased both frequencies and amplitudes in GluDTR islets but only to 64±4 % of WT islets. We also tested for the contribution of GLP-1 receptors in WT islets using a relatively stable antagonist, Exendin 9-39 to avoid tissue degradation of GLP-1 and the potential generation of antagonist intermediates (40). The presence of exendin 9-39 considerably reduced amplitude and frequency mainly throughout the 2^nd^ phase (Fig. S1) suggesting a role for GLP-1 receptors in the electrical activation of islets by glucose and amino acids.

**Figure 3:**
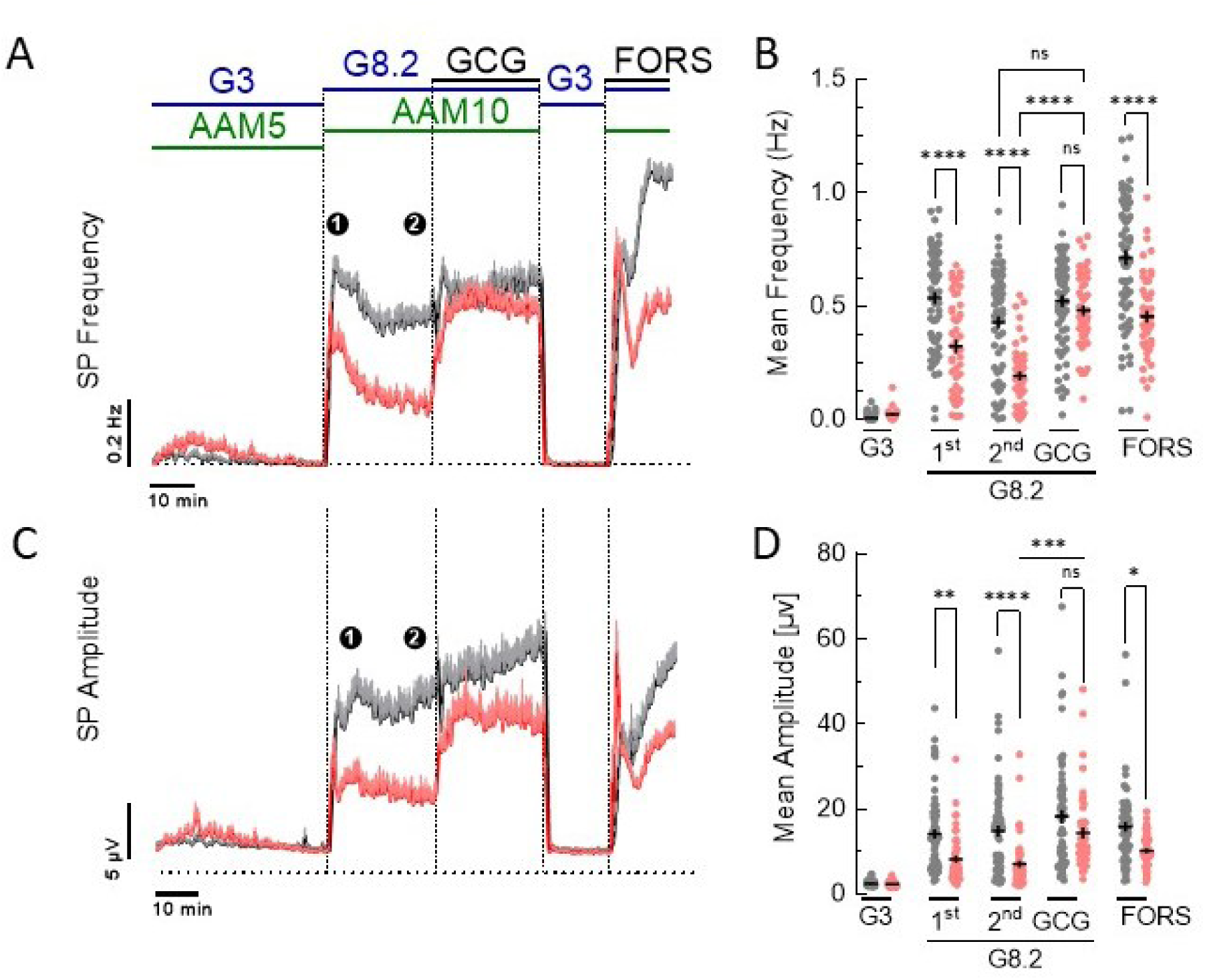
Effect of glucagon and forskolin on electrical activity stimulated by glucose and amino acids in WT and GluDTR islets. (**A)** Effect of glucose (3 mM, G3; 8.2 mM G8.2), amino acids (5 mM, AAM5; 10 mM AAM10), glucagon (GCG, 1 nM) or forskolin (FORS, 1 µM) on mean slow potential frequencies (±SEM). 1^st^ and 2^nd^ phases are indicated (❶,❷). (**B**) Statistics of mean frequencies. N=4, n=49-68. (**C)** Effect of glucose, amino acids, glucagon or forskolin (concentrations as in A) on mean slow potential amplitudes (±SEM). (❶,❷). (**D**) Statistics of mean amplitudes. 2-way ANOVA and Tukey; *,**,***,****, 2p<0.05, <0.01, <0.001, <0.0001.

Subsequently we measured a distal outcome of transmembrane ion fluxes, that is insulin secretion from islets (Fig. 4). At low glucose levels (G3) no difference was observed between WT and GluDTR islets. G8 stimulated secretion from WT islets about 5-fold but only 2.3-fold in GluDTR islets and the addition of AAM increased secretion in WT islets 10-fold versus only three-fold in GluDTR islets (Fig. 4A). The addition of glucagon largely restored insulin secretion in GluDTR islets to WT levels regardless of the absence or presence of AAM. Similarly, GLP-1 increased glucose-induced insulin secretion in WT and GluDTR islets in the absence or presence of AAM. However, in contrast to glucagon, GLP-1 restore secretion in GluDTR only to about 70% of levels observed in WT islets (Fig. 4A). We also examined the effect of molecules that act downstream from receptors (Fig. 4B). Forskolin (1 µM), a direct activator of adenylyl cyclases, induced a robust enhancement of insulin secretion at G8.2 in WT islets in the absence or presence of AAM whereas only a far smaller change was observed in GluDTR islets (17.0±4.3% of WT at G8.2, 46.6±10.2 at G8.2 with AAM). Increasing the forskolin concentration to 10 µM stimulated insulin secretion in GluDTR islets at G8.2 only to 17.0±4.9% of WT islets (N=8), the addition of the phosphodiesterase inhibitor IBMX (0.1 mM) to forskolin at G8.2 stimulated GluDTR islets to 53.3±5.1% of WT islets (N=3) and forskolin at G16.7 stimulated GluDTR islets to 37.8±3.9% of WT islets. Likewise, membrane depolarization by KCl (at G3) produced a considerable increase in insulin secretion but only in WT islets. Finally, mastoparan, a direct stimulator of exocytosis (41), increased insulin secretion to similar extent in WT and GluDTR islets. Collectively these data indicate a major role for glucagon as well as for cAMP-dependent and -independent pathways in the observed phenotypes.

**Figure 4:**
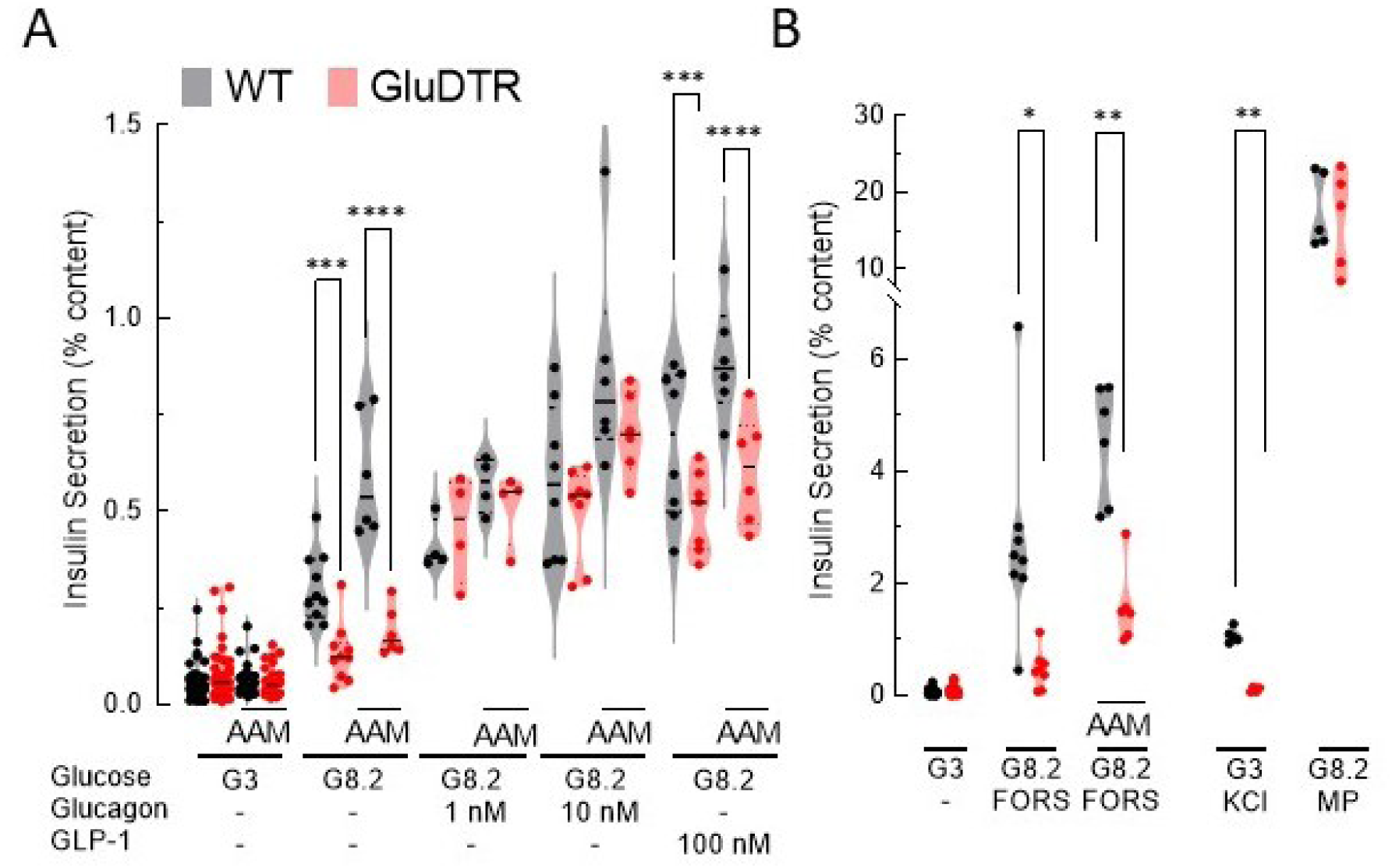
Insulin secretion from WT or GluDTR islets. **(A)** During secretion assays, islets were incubated with 3 or 8.2 mM glucose (G3, G8.2) in the absence or presence of amino acids (AAM, 5 mM at G3; 10 mM at G8.2), and in the absence or presence of glucagon or GLP-1. N=3-6, n=12-77. (**B)** Islets were incubated in the absence or presence of amino acids (AA, 10 mM) with 3 or 8.2 mM glucose (G3, G8.2) and in the absence or presence of KCl (24 mM), forskolin (FORS, 1 µM) or mastoparan (MP, 30 µM). N=3-6, n=12- Mann-Whitney (Benjamini, Krieger, and Yekutieli); *, **, ***, ****, 2p<0.05, <0.01, <0.001, <0.0001.

### α-cells enhance β-cell synchrony and cluster size

Next, we set out to decipher the dynamic modulation of the intra-islet β-cell networks by the presence/absence of α-cells during physiological stimulation by glucose and amino acids. To this end we used microelectrode arrays with high-density coverage by electrodes (HD-MEA) to monitor regional changes in electrical activity within islets. As shown in Fig. 5A, electrodes are spaced 30 µm apart corresponding to about 2 to 3 β-cell diameters. SPs can be correlated to establish correlation matrices and presence of clusters (>1 correlated electrode, Fig. 5A). Pairwise identifications of phase shifts allow to establish the order of activation, leader regions, extent of connectivities within islets and speed of propagation (Fig. S2). Original recordings are given in Fig. S3 and show again the large reduction of electrical activity in GluDTR islets as compared to WT islets in response to glucose and amino acids. Since differences in electrical activity between WT and GluDTR islets were most marked in the presence of amino acids, we have concentrated our analysis on the effects of glucose in the presence of AAM. As shown in Fig. 5B and C, synchrony in WT islets increased considerably from G3/5mM AAM to G6/10 mM AAM and G8.2/10 mM AAM. Synchrony in GluDTR islets at G3/5mM AAM was widely scattered but cluster remained small and at G8.2/10 mM AAM a strong difference was evident between WT and GluDTR islets. Figure 5D gives an example of the distribution of synchronous electrodes and the presence of multiple clusters of different sizes. Increasing glucose from G3 to G6 and G8.2 with concomitant increase in amino acids considerably increased the relative size of clusters in WT islets from 0.1 to 0.6 and 0.8 whereas in GluDTR islets only an increase from 0.1 to 0.2 and 0.4 was observed. This was also apparent when grouping clusters according to their sizes and looking at their distribution (Fig. S4). Whereas cluster sizes of WT islets increased already at G6/10 mM AAM, this was only apparent in GluDTR islets at G8.2/10 mM AAM. Moreover, GluDTR islets never formed very large clusters. Thus, cluster sizes at stimulatory glucose and amino acid concentrations were significantly smaller in GluDTR islets. We also performed a correlation analysis of cluster sizes, synchrony, mean frequencies and amplitudes at G3, G6 and G8.2 in the presence of AAM (Fig. S5). Synchrony correlated significantly with SP amplitudes and confirms that amplitudes may serve as a reliable surrogate for β-cell coupling.

**Figure 5:**
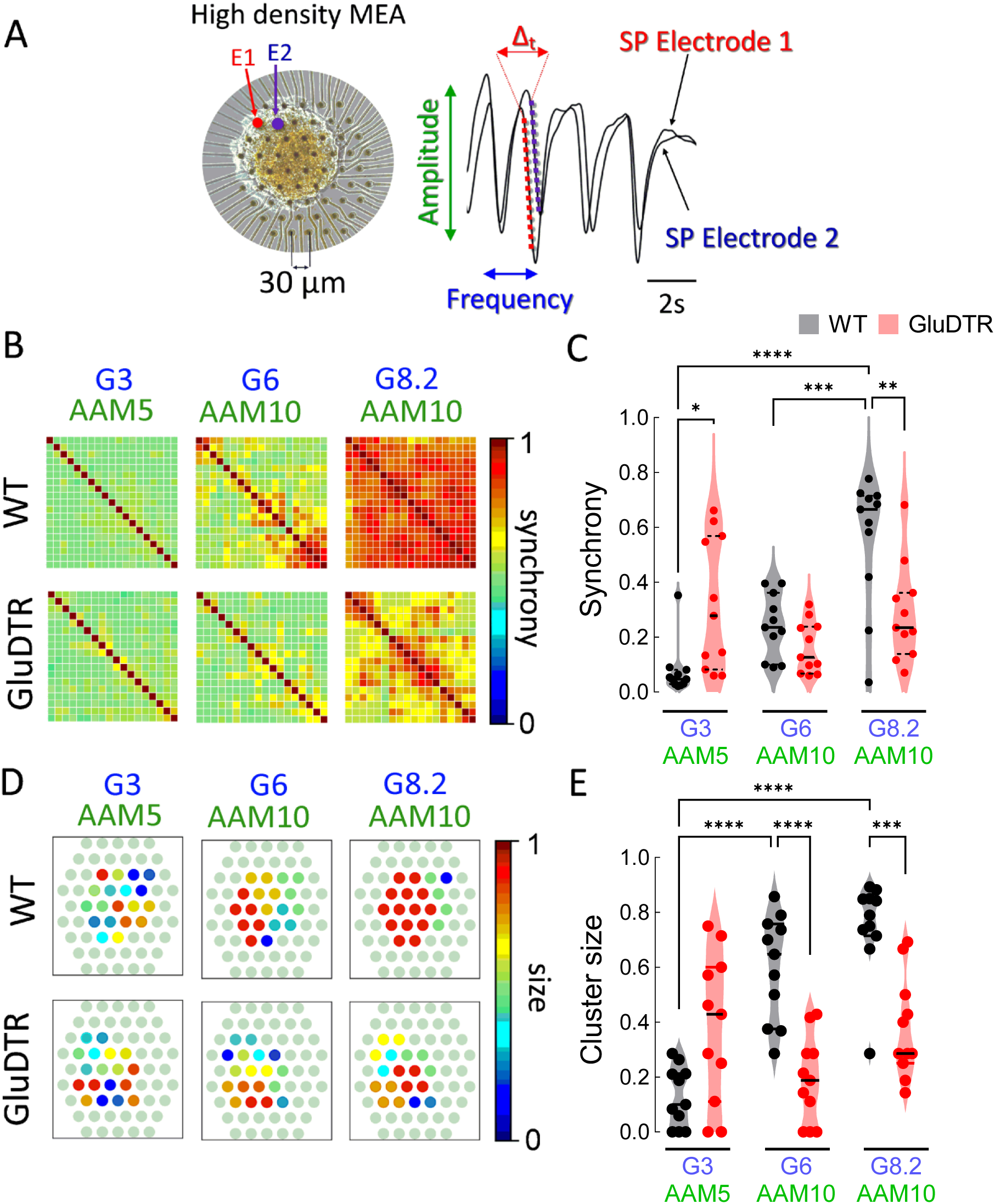
Effect of glucose and amino acids on synchrony and clusters in WT and GluDTR islets. (**A)** Image of an islet on a high-density MEA and example of two traces obtained from two neighboring electrodes. The electrodes (Ø 10 µm) are spaced 30 µm apart (electrode borders). (**B**) Correlation plot i.e. synchrony within islets exposed to different concentrations of glucose (3, 6 or 8.2 mM; G3, G6, G8.2) and amino acids (5 or 10 mM, AA5, AA10). Each tile represents an electrode pair. (**C**) Statistics of B. (**D**) R Spatial representation of cluster size. Representative outputs of cluster sizes in the different conditions are given. The normalized size of different clusters is indicated by color codes (normalization to the size of each islet). (**E**) Statistics on cluster sizes. N=3 (animals), n=11 islets for each condition. 2-way ANOVA and Tukey; *, **, ***, ****, 2p<0.05, <0.01, <0.001, <0.0001.

### α-cells increase β-cell cluster leader regions and signal speed, but not their life span

We subsequently addressed the spatial origin of electrical activity, which we termed “leader region”, and the spatial evolution of islet activation (Fig. 6). To that end the 20 min stimulation period (G8.2/10 mM AAM) was divided in three consecutive periods and the relative ranking of a given electrode as leading region during that period was determined and presented with a color code in a pie chart. Figure 6A shows an example of a single WT or GluDTR islet. Only a limited number of electrodes were marked with red segments (indicating first leader regions) in most of their pie charts, although clearly distinct and widely separated regions could take the lead. While the number of active regions is fairly reduced in GluDTR islets, again red segments were concentrated in a few regions. As there was little evidence for clear cut differences between the three time periods, we pooled them for a global analysis (Fig. 6 B-K). To indicate the relative frequency at which an electrode was present, the most prevalent was termed first leading region (LR^1st^), the second or third most prevalent leading region termed LR^2nd^ and LR^3rd^. Also given are regions that occur at an even lower frequency (“other”) as well as the absence of any detectable leading region. As there was a large difference between WT and GluDTR islets in regard of the presence of any leading region, we also normalized the data by excluding the occurrence of no discernible leading region (Fig. 6 B).

**Figure 6:**
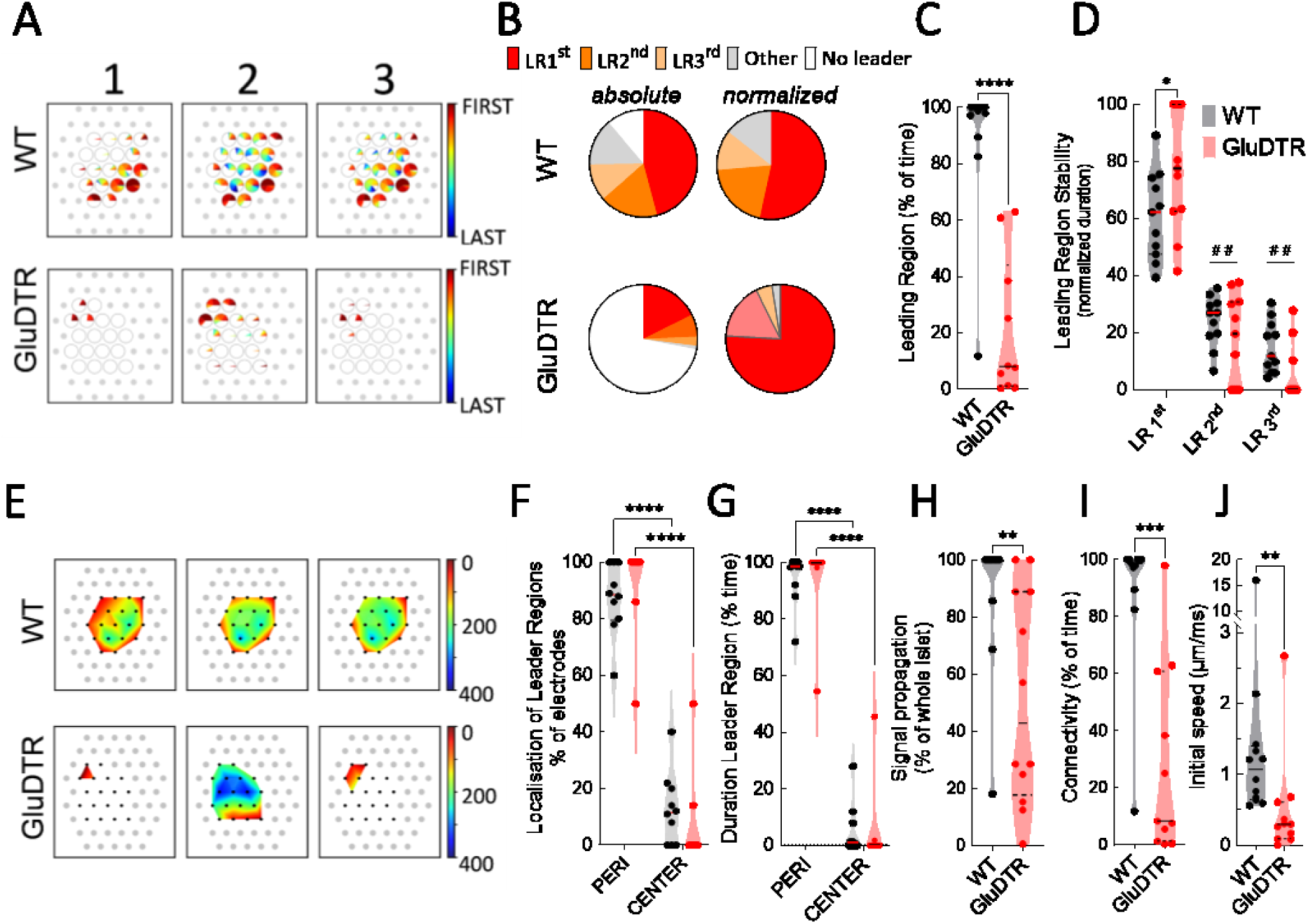
Properties of leader regions in WT and GluDTR islets. All data were obtained during stimulation with 8.2 mM glucose and amino acids. (**A)** Example for the order of appearance of activity (first to last) and relative distribution of electrodes (“regions”) as pie charts under one islet, either WT or GluDTR. Given is the distribution during three periods (1, first 300 s of stimulation; 2, following 300 s; 3, last 300 seconds) during by 8.2 mM glucose in the presence of amino acids. Open or filled circles, electrodes covered by an islet; grey dots, non-covered electrodes. (**B**) Absolute and normalized frequencies of same electrode positions as leader region frequencies (1^st^, most often; 2^nd^ and 3^rd^ most often; others, 4^th^ and more most often) for all experiments (N=3 animals, n=11 islets for each condition). The normalized pie chart excludes the absence of leader regions. (**C**) Statistics of the detection of leading regions in WT or GluDTR during the three periods. WT, black; GluDTR, red. (**D**) Appearance of the same electrode throughout phase 1, 2 and 3 as most common leading region (LR 1^st^), second or third most common leading region (LR 2^nd^, LR 3^rd^). **(E)** Propagation time of activation of one islet (example; WT or GluDTR) at a color-coded time scale from 0 to 400 ms (right Y axis). Black dots, electrodes covered by islet; grey dots, other electrodes. (**F**) localization of leader regions in the periphery (PERI) or center (CENTER) of islets. (**G**) duration of the leader regions in the periphery or center of the islets. (**H**) Extent of spatial signal propagation within islets. (**I**) Duration of connectivity between regions for all experiments in combined period 1, 2 and 3, given as percent of time. (**J**) initial speed of signal propagation (between 1^st^ leading electrode and following electrode). N=3 (animals), n=11 islets for each condition. C, F-H, Mann Whitney; D, I, J, 2-way ANOVA and Tukey; **, ***, ****, 2p<0.01, <0.001, <0.0001; ^##^, 2p<0.01 as compared to corresponding LR1^st^.

In WT islets the same region was the leading one for over half of the time and during 85% of time the leading region is defined by just three different electrodes which may, however, be widely spaced apart. The situation is similar in GluDTR islets where in 2/3 of the events the connectivity started in the same region. We also performed statistics on the stability of the most frequent leading region as well as the second and third most often occurring regions (Fig. 6 C). The life span of leading regions, i.e. stability, was significantly different in both types of islets between the most frequent LRs and the less frequently occurring regions, LR^2nd^ and LR^3rd^ (Fig. 6 D). Notably, the stability did not vary between WT and GluDTR: a given LR^1st^ lasted for about 20 to 30 s in WT as well as GluDTR islets, with maximal durations of more than 100 s, and decreased to about 10 s for LR3^rd^ (Fig. S6A, B). The mean distance between successive leader regions in WT islets was 88±9 µm, which is almost 3 electrodes apart, and decreased significantly to 62±7 µm in GluDTR islets (Fig. S6 C).

Next, we examined the propagation of the activation throughout the islets. As given in the examples here (Fig. 6 E), activation in most instances started from the periphery in WT as well as in GluDTR islets and peripheral leader regions were more stable than central ones (Fig. 6 F, G). In WT islets this most often led to an activation throughout the entire islet (Fig. 6 H) whereas a high variability was observed in GluDTR islets. Moreover, in WT islets a network (termed connectivity) was maintained during most of the time whereas this was the case for only about 10% of the time in GluDTR islets (Fig. 6 I). The initial speed of activation, defined as time lag between leading and following region, was significantly slowed down from 2.3±1.2 µm/s in WT islets to 0.5±0.2 µm/s in GluDTR islets (Fig. 6 J).

## Discussion

During the last years the view on α-cells has evolved from a mainly counterregulatory task in glucose homeostasis to more nuanced intra-islet cooperativity in handling of the physiological nutrients. However, a detailed view on resulting β-cell activity and networks is still lacking especially in electrophysiological terms. Our analysis of GluDTR mice confirms that α-cells are pivotal for nutrient homeostasis including amino-acids in-vivo and islet activity in-vitro in mice. Although the effect of amino acids on glucagon secretion was known for long time, previously in-vivo experiments have mainly been conducted using only one or some selected amino acids which may not represent a physiological situation due to the wide differences in potency of individual amino acids on glucagon as well as insulin secretion (41).

The use of extracellular electrophysiology now defines precisely the effect of amino-acids on β-cells and its mediation by α-cells. The activation of β-cells in islets proceeds by two phases. Whereas the short-lasting first phase is characterized by a high activity with little β/β -cell coupling, the second phase exhibits a lower frequency of activity but a higher degree of coupling reflected by increased amplitudes and synchrony (33). In wild-type islets, the amino acid mix (AAM) increased the mean slow potential frequency in the first and second phase at 6 and 8.2 mM glucose. The mean amplitude changed only slightly with AAM at 8.2 mM glucose, likely because glucose alone already reached almost maximal stimulation (36). Notably, GluDTR islets consistently showed lower mean frequency and amplitude of the second phase when amino acids were present. The amplitude of slow potentials varies with the degree of β-cell coupling (33), and the reduced amplitudes observed are consistent with the observed smaller cluster size and lesser coupling in GluDTR islet β-cells. In the absence of amino acids and sole presence of glucose, we did not observe a significant difference in mean electrical activities between wild-type and GluDTR islets, despite the fact that GluDTR islets secreted considerably less insulin under those conditions as also has been reported in other studies using the GluDTR model or β-cell specific knock-out of the glucagon or GLP-1 receptor (10, 12, 24, 42). The discrepancy between reduced insulin secretion in-vitro and the preserved glucose tolerance in-vivo is probably due to increased GLP-1 levels in-vivo in GluDTR mice (24). The difference observed in-vitro upon stimulation by glucose only between unaltered electrical activity and reduced secretion suggests that α-cell hormones may alter signal transduction distal from ion fluxes in β-cells. It is indeed known that glucagon or GLP-1 receptors can activate not only adenylate cyclases but also phospholipases and phosphatidylinositol 3-kinase which all alter insulin secretion independently from ion fluxes (43–46).

We were not able to restore electrical activity or secretion by forskolin, a general activator of adenylyl cyclases, as had been reported by others measuring secretion in the GluDTR model in the same genetic background (24) or in glucagon receptor deficient mice islets (10). We have no ready explanation for the difference in forskolin effects but are intrigued by the fact that only forskolin but not the general phosphodiesterase inhibitor IBMX had been reported to restore secretion in glucagon receptor knock-out mice (10). This points towards the requirement of extremely high levels of cAMP as low doses of IBMX already increase cAMP in β-cells to a larger degree than glucose itself (47). The final steps in secretion were not altered between wild-type and GluDTR islets as mastoparan, a direct stimulator of exocytosis (40), caused comparable effects.

Our recovery experiments suggest that glucagon is the major α-cell derived mediator. Results different from ours have been reported (24) and may be due to a higher concentrations of GLP-1, as we used only 1 nM and not 10 nM since the levels of α-cell derived GLP-1 have been reported to be extremely low (11–12, 48). Although the precise local concentration in the vicinity of β-cells is unknown, already 1 nM is far above the threshold of exogeneous GLP-1 concentrations required to stimulate β-cells (17, 33, 36). Notably, the differences in transmembrane ion fluxes and in insulin secretion observed in GluDTR islets were recovered by glucagon as reported previously for hormone secretion (24). In contrast, GLP-1 was unable to fully restore insulin secretion whereas the GLP-1 receptor antagonist exendin 9 reduced activity in wild-type islets. Eventually both receptors, for GLP-1 and glucagon, have to be activated to recover wild type-like activity as glucagon is known to act via these two receptors (12, 18, 49). Collectively this pin-points to the pivotal role of glucagon acting via both, glucagon and GLP-1 receptors, as suggested by other studies (13).

The use of MEAs and high-density MEAs has advantages and limitations. Electrophysiology offers a far greater time resolution than optical means where frequency of data acquisition is generally limited to about 0.5 or 1 Hz for a few cell layers as compared to the kHz range in our case. Note that most recent work was capable to resolve events such as action potentials (50) for a short time span or to image entire islets at 2 Hz with cellular resolution for an entire islet (51). In contrast to imaging, even high density MEAs lack single-cell resolution and we can only determine regions. Moreover, slow potentials represent summation signals from more than one single cell (33) which also precludes determination of single cell origin but their analysis considerably reduces noise and increases robustness. Actually, most of the imaging leading to the concept of hubs and leaders was done by recording only one or two layers of the islet surface due to technical constraints by acquisition speed and probe penetrance (52–53). Although MEA recordings cannot distinguish between ion species, quantitatively most of the observed changes in field potentials are carried by Ca^2+^-mediated depolarization in β-cells (29) and are therefore mostly representing the same ion flux as observed by calcium imaging. We would, however, like to point out that slow potentials are far more rapid phenomena of much shorter duration and occurring at higher frequency than the Ca^2+^ hub or leader cell activities as well as Ca^2+^ waves that have been used for network analysis by imaging. Whereas SPs last around 1 second (33, 54), reported leader or hub activities are in the range of 10 seconds and more and waves last several minutes (52–53, 55). Most recently a very elegant approach has been reported using multiple MEA meshes with a lower resolution (electrodes 60 µm apart, electrode center to center) but which permit recording from intra-islet sites (56). However, this powerful approach requires islet dissociation and reaggregation into pseudo-islets, a procedure that may change native topology. Another limitation of our study is the absence of vascularization especially as the intra-islet blood flow is often thought to proceed vectorially from β- to α-cells and thus in a native settings β-cells would receive little input from α-cells especially in the rodent islet. This issue is a matter of discussion as different vascular perfusion patterns have been described and α-cells are not only present in the mantle of rodent islets (57–58).

Islet β-cells are functionally heterogeneous (59–60) and achieve physiological effects by coupling and networks driven by gap junctions and cell intrinsic dynamics (61–62). Previous work based on calcium imaging has pointed to the presence of a stable number of few hub or leader cells with distinct expression profiles which initiate wave-like islet activity (52–53, 62) although a more dynamic and nuanced behavior of leaders and first responders has been reported more recently (51, 63–64). The spatial organization of electrical activity in WT islets indicated a preponderance of a defined region as the starting point for cluster formation. However, despite this marked occurrence in a given region, the leading region was not at all limited to one area but intermittently other distant regions of the islet took the lead. Even though our data contain a limited spatial resolution, these observations are not easily reconcilable with the presence of defined and stable leader or first responders at least under our more physiological conditions, i.e. the stimulation by glucose and amino acids.

Remarkably SP activity often started at the border of the islets, similar to what has been reported for Ca^2+^-waves in slices or whole islet imaging (51, 65–66). In rodent islets, α-cells are more often present in the periphery and the presence of preproglucagon-derived peptides may lower the activation threshold. However, we do not think that this predilection for the periphery may be brought upon by the special α-/β-cell topography as this signal topology was also apparent here in GluDTR islets which renders this mechanism unlikely. Coupling among β-cells does not only provide efficient means for concerted activation but will also increase the activation threshold thus dampening β-cell activity (67–69). As at the periphery of the islets β-cells have fewer neighboring cells for coupling, their activation threshold may be lower.

The analysis of synchrony and cluster size revealed considerable smaller clusters in GluDTR islets and an overall less organized functional activity at increased glucose and amino acids which is reflected also in diminished insulin secretion. Indeed, proglucagon-derived peptides are known to reduce the glucose-dependent activation threshold of islet β-cells (18, 70–71) and their absence may explain the lower fraction of active islet β-cells in GluDTR islets as well as reduced initial speed and signal propagation. Notably, the life span of first, second and third leader regions (LR^1st^, LR^2nd^, LR^3d^) was not different in GluDTR. Thus, frequency of activation, speed and coupling is regulated by α-cells. In GluDTR islets probably only those leader regions with the lowest activation threshold respond and the presence of α-cell hormones, such as in WT islets, lowers the general activation threshold and increases the total number of leader regions, propagation speed and extension.

The reduced insulin-secretion in-vivo upon glucose and amino acid tolerance tests observed here provides strong evidence that α-cells are not only required upon metabolic stress as thought previously (10, 21–23), but also under more physiological conditions. This specific observation combined with the electrophysiological characterization allows us to propose that the α-β interaction via proglucagon peptides is required for proper islet activity via the functional properties of the β-cell network. Elegant in-depth characterization of mice harboring only β-cells has demonstrated that these animals have an improved glucose tolerance and highly adequate β-cell function even under metabolic stress (25). It will be interesting to test whether a more equilibrated exposure to nutrients, including also amino acids, may reveal subtle differences and potentially altered functional β-cell networks in this model. Detailed knowledge of the dynamic role of α-cells may also help to improve currently used algorithms that control therapeutic insulin delivery in diabetes and are mainly based on glucose sensing (1, 72–73).

## Materials and methods

### Materials

Glucagon, forskolin and mastoparan were purchased from Sigma-Aldrich (St. Louis, MO, USA), exendin9-39 and GLP-1 from Bachem (Bubendorf, Switzerland).

### Animals and islets

GluDTR and corresponding wild-type mice (in C57BL/6N genetic background) were kindly provided by Dr Marc Donath (21, 24). Diptheria toxin (D0564; Sigma; dissolved in 0.9% w/v NaCl) was injected in 5- to 6-week old WT and GluDTR animals in three i.p. injections of 500 ng each on days 1, 3, and 5 as reported before (21, 24). Mice were used for experiments 6 to 20 weeks after injections. For genotyping the following primers were used: G1-52S: 5’- GAG AAA TTT ATA TTG TCA GCG -3’; DTR reverse 4: 5’- CTT CAG CAC CAC CGA CGG C -3’ resulting in a ∼0.8 kb PCR transcript. PCR was performed as published by Traub et al. (24). Mice were sacrificed by cervical dislocation according to University of Bordeaux ethics committee guidelines (authorization number #2087-2019052917497896). Islets were obtained by enzymatic digestion and handpicking (31, 33–34).

### Study approval

All animal experiments were reviewed by the Bordeaux University Ethical Committee with the corresponding government authorization (PA number) #2087-2019052917497896.

### In-vivo characterizations

Glucose (2 g or 3/kg body weight) and amino acids (1g/kg body weight) or pyruvate (2 g/kg body weight) were administered by i.p. injection after 6 h of fasting (8:00–14:00) (34). Blood samples were obtained by tail tip bleeding and assayed using a glucometer (Freestyle; Abbott) and collected on hematocrit capillary tube (Hirschmann™ 9100275). Blood samples collected at different time points were transferred to low-binding Eppendorf tubes (Sorenson 11300), centrifuged 15 min at 1800g at 4°C and the plasma transferred in a new tube and frozen at 20°C

### Insulin secretion

Insulin secretion was determined under static conditions. After 4 days of culture in RPMI medium, to allow recovery from isolation-induced stress and in line with the duration of islet culture on MEAs, 40 islets were placed on small filters inserted into the wells of a 48-well plate (PluriStrainer Mini 40 µm, Dutscher) containing 500 μL of EPHYS solution (composition see below, Electrophysiology). Experiments were conducted in this buffer to ensure comparability with MEA experiments. The filters allowed easy and rapid transfer of islets from one well to another. All incubations were performed at 37°C and 5% CO_2_ for 30 min. An initial incubation served to wash the islets from the culture medium, followed by basal (3 mM glucose) and stimulated conditions. For stimulations in the presence of amino acids, the same islets and wells were kept for 3h in culture medium prior to washes and incubations as above. Finally, the islets were placed in ethanolic acid at −20°C for at least 12 h. All solutions were collected, centrifuged at 800 g for 5 min and 300 µL of the supernatant was stored at −20°C. Islets were taken up in acid-ethanol and kept at −80°C until analysis. Insulin concentrations were determined by ELISA (Mercodia mouse insulin; 10-1247-01) as was Glucagon (Mercodia Glucagon 10-1281-01).

### Electrophysiology

Islets were seeded on MEAs coated with Matrigel (2% v/v) (BD Biosciences, San Diego, CAand cultured at 37°C (5% CO2, 0.9% relative humidity) in RPMI medium (11 mmol/L glucose; Thermo Fisher Scientific, Waltham, MA) and medium was changed every other day as described (33–34, 74). Experiments were performed at 37°C in EPHYS buffer containing (in mM) NaCl 135, KCl 4.8, MgCl2 1.2, CaCl2 1.2, HEPES 10 and glucose and amino acids as indicated (pH 7.4 adjusted with NaOH). The physiological amino acid mix AAM10 was according to Zhu et al. (26) and contained: Ala 0.88, Arg 0.38, Asp 0.076, Cit 0.19, Glut 0.24, Gly 0.6, His 0.15, Ile 0.19, Leu 0.32, Lys 0.74, Met 0.1, Orn 1.4, Phe 0.16, Pro 0.7, Ser 1.14, Thre 0.54, Trp 0.15, Val 0.4 and Glut 2 (in mM). AAM 5 contained half the amount of each amino acid. Extracellular recordings were performed on MEA placed in a MEA recording system (USB-MEA60-Inv-System-E amplifier (MCS; gain: 1,200, Multi-Channel Systems GmbH [MCS], Reutlingen, Germany) controlled by MC_Rack software (v4.6.2, MCS). Recordings of different intra-islet regions were performed using high-density MEAs (HD-MEAs) (60HexaMEA40/10iR-ITO-gr, 59 TiN electrodes, Ø 10 mm, 30 mm between electrodes, border to border) that were continuously perfused at 0.5 mL/min (Reglo ICC; Ismatec, Glattbrugg, Switzerland). Extracellular field potentials were acquired at 10 kHz, amplified, and band-pass filtered at 0.1–3,000 Hz (31–33, 36–37, 74). Images of islets on MEAs were taken before and after each experiment to localize electrodes covered with islets. Electrophysiological data were analyzed with MC_Rack software. Slow potentials (SPs) were isolated using a 0.1-2 Hz bandpass filter and frequencies were determined using the threshold module of MC_Rack with a dead time (minimum time between two events) of 300 ms. The peak-to-peak amplitude module of MC_Rack was used to determine SP amplitudes.

### Analysis of HD-MEA recordings

Synchrony measurements, phase measurements, clustering, and order of activation were computed with custom Python software, using libraries numpy, pandas, and scipy. Libraries matplotlib and seaborn were also utilized for data visualization. Electrophysiological data were filtered using 0.2-2.0 Hz Bessel filters and resampled at 100 Hz. Synchrony between signals was measured using pairwise Pearson correlation between signals from all electrodes (Figure S1). From the resulting correlation matrix, clusters of synchronized electrodes were identified through hierarchical clustering using UPGMA (unweighted pair group method with arithmetic mean). The clustering threshold was set to 70% of the maximum distance between signals as calculated in the linkage matrix. Lag between signals was measured by computing the cross-correlation between signal pairs and retrieving the lag at maximum correlation for each pair. Due to the 100 Hz sample rate, time resolution for phase measurements was 10 ms. All lag values that yielded a maximum correlation below 0.7 were discarded. To monitor changes over time, signal lag was computed in a 30 s sliding window, with steps of 7.5 s (75% overlap). The order of activation of electrodes was deduced from the windowed lag measurements, by identifying the leader in each window (signal of earliest lag) and sorting the others by ascending phase, relative to the leader. For analysis periphery versus central, electrodes were classified based on their relative position with respect to the islet outline. The peripheral region was defined as the first continuous outer line of electrodes covering the islet boundary. All remaining electrodes located within this peripheral layer were classified as central. This procedure was repeated for each individual islet to account for variability in islet size and position.

### Statistics

Graphics, quantifications, and statistics were performed with Prism software (v7; GraphPad, La Jolla, CA). Data are presented as means and SEM. The minimal value of mean SP frequency after the first peak (corresponding to the nadir) was taken as the limit between phases. Gaussian distributions were tested by Shapiro-Wilk test and statistical tests performed as indicated in the figure legends.

## Acknowledgments

We gratefully acknowledge the help by Dr Marc Donath providing us with the relevant mouse strains and useful information.

## Data availability

The data that support the findings of this study are available from the corresponding author upon reasonable request.

## Author contributions

ML: conceptualization, formal analysis, investigation, methodology, visualization, data curation, writing – original draft, writing – review & editing. MJ: conceptualization, formal analysis, investigation, data curation, writing – review & editing. AP: formal analysis, investigation, data curation, writing – original draft, writing – review & editing. KLF: formal analysis, investigation, data curation. JG: methodology, investigation. SR: supervision, review & editing; funding acquisition. MR: conceptualization, methodology, formal analysis, investigation; supervision; writing – original draft, writing – review & editing; funding acquisition. JL: conceptualization, formal analysis, methodology, visualization, data curation, writing – original draft, writing – review & editing; funding acquisition.

## Supplementary Materials

**Fig. S1:**
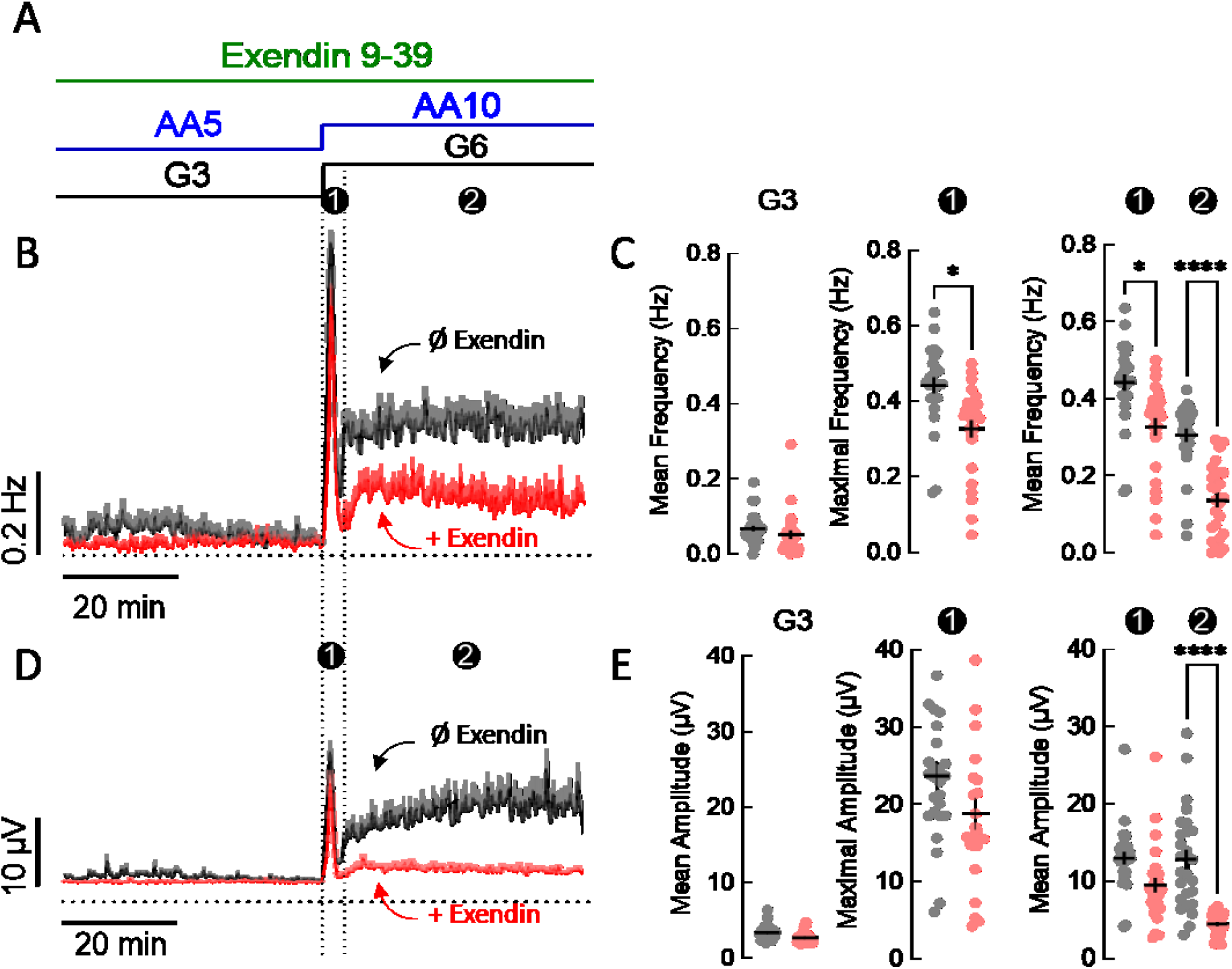
Effect of the GLP-1 antagonist Exendin 9-39 on glucose and amino-acid evoked electrical activity in WT islets. (**A)** Incubation scheme, glucose (3 mM, G3; 6 mM G6), amino acids (5 mM, AA5; 10 mM AA 10) and Exendin 9-39 (1 nM). Black, no exendin 9-39; red, in the presence of exendin 9-39. (**B)** Mean slow potential frequencies (±SEM). 1^st^ and 2^nd^ phases are indicated (❶,❷). N= 2, n=26. (**C)** Statistics of frequencies. (**D, E**) Mean slow potential amplitudes (±SEM) and statistics. 2-way ANOVA and Kruskal; *, **, ***, ****, 2p<0.05, <0.01, <0.001, <0.0001.

**Fig. S2:**
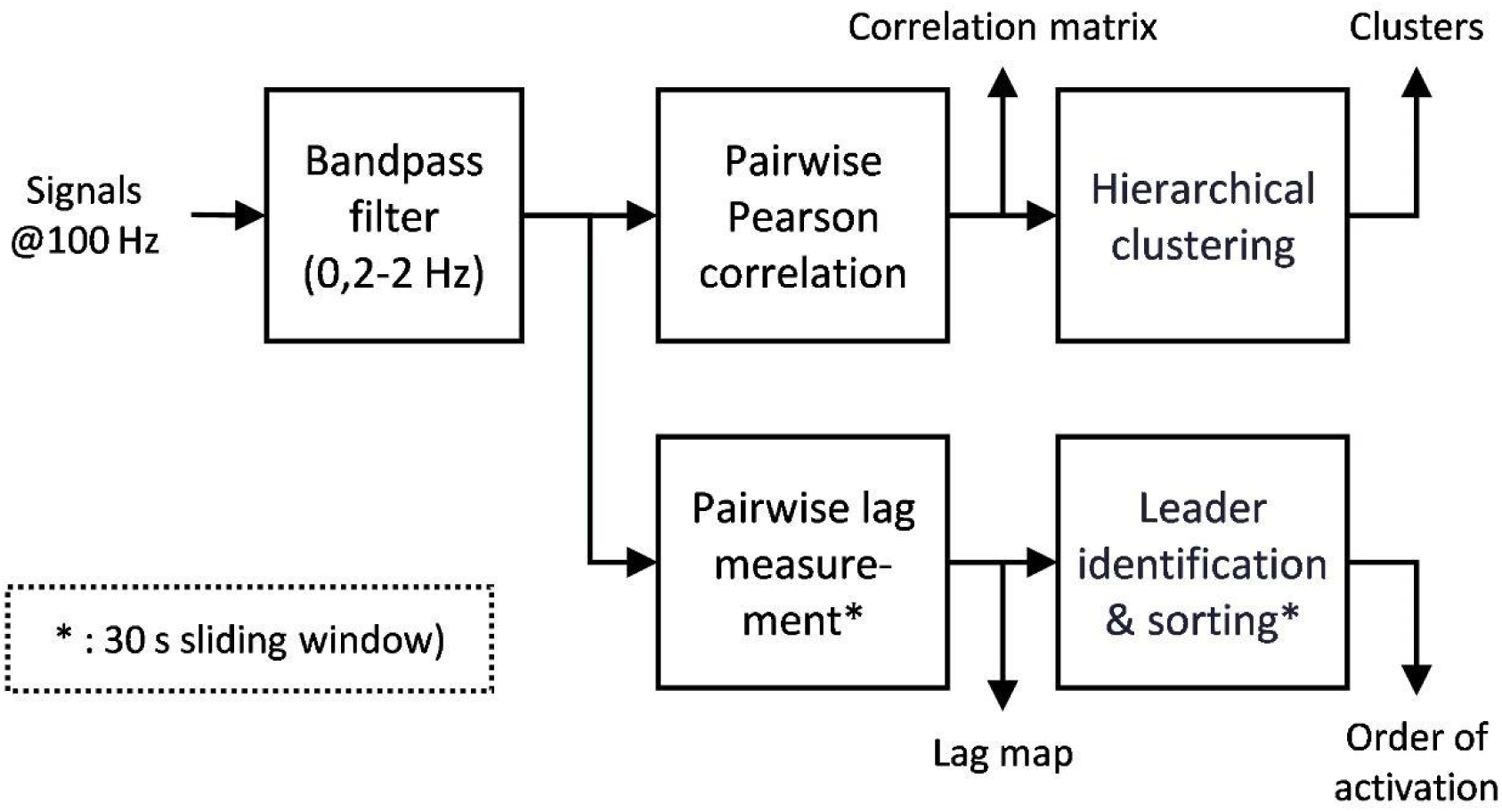
Scheme of data analysis of HD MEA data.

**Fig. S3:**
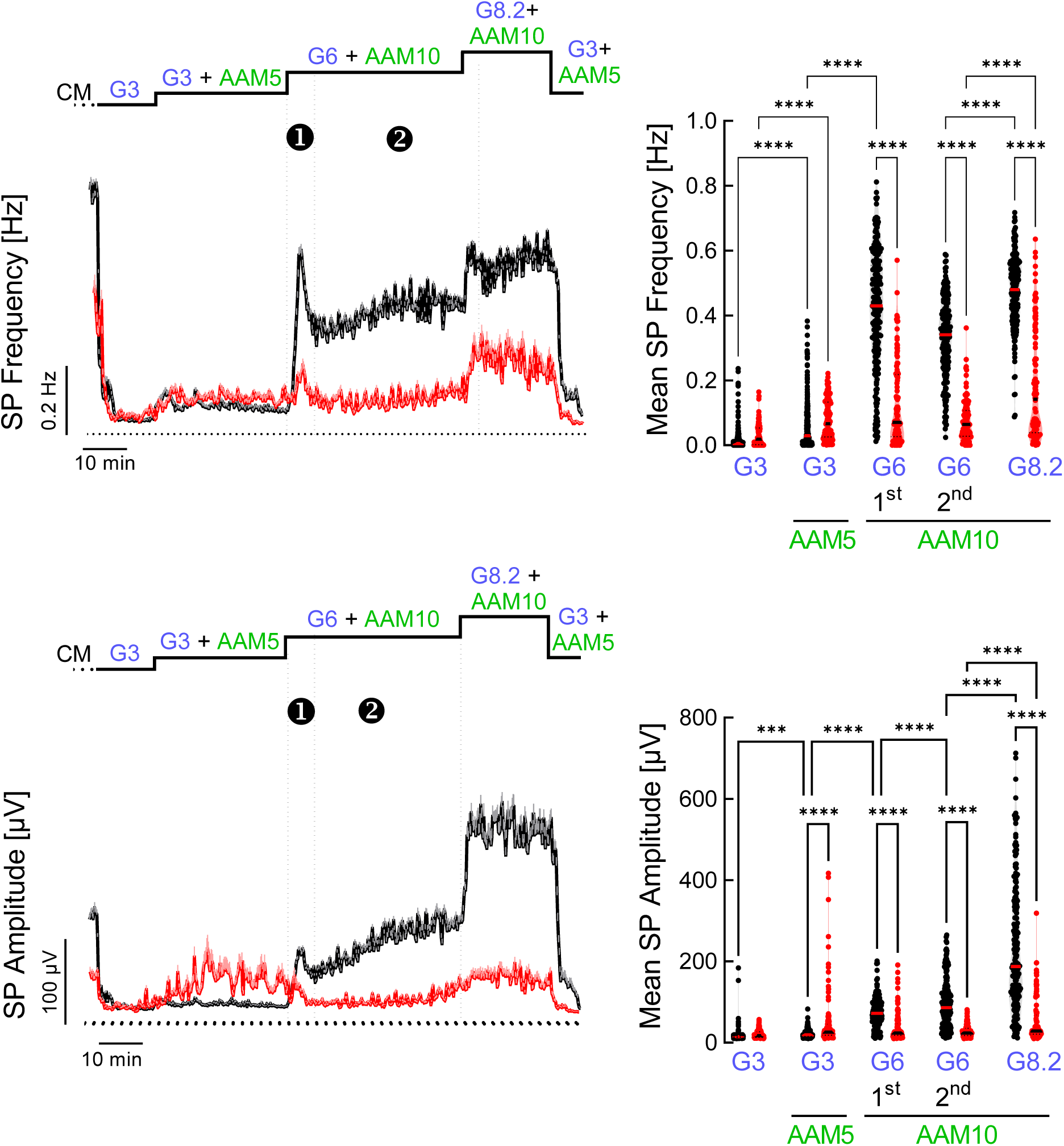
Recordings of WT and GluDTR islets on high-density MEAs. (**A)** Effect of glucose (3 mM, G3; 8 mM G8) and amino acids (5 mM, AA5; 10 mM AA 10) on mean slow potential frequencies (±SEM). 1^st^ and 2^nd^ phases are indicated (❶,❷). (**B**) Statistics of mean frequencies. (**C)** Effect of glucose and amino acids on mean slow potential amplitudes (±SEM). (❶,❷). (**D**) Statistics of mean amplitudes. 3 animals and 11 islets total; 2-way ANOVA and Tukey; ***, ****, 2p <0.001, <0.0001.

**Fig. S4:**
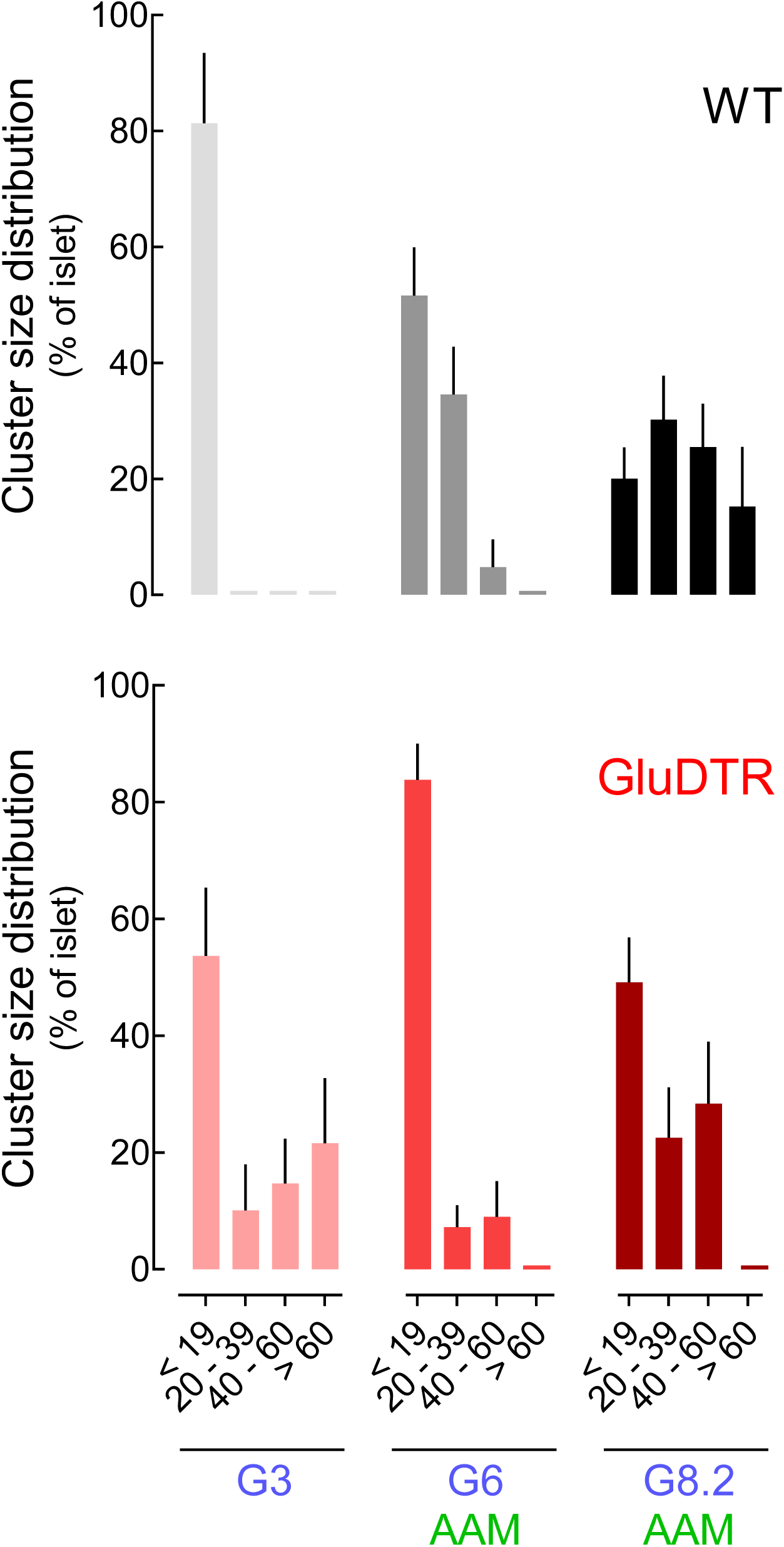
Histogram of cluster size distribution. Islets were exposed to 3 mM glucose (G3), 6 mM glucose and amino acids (G6 AAM) or 8.2 mM glucose and amino acids (G8.2 AAM) in WT or GluDTR islets. (**A**) WT; (**B**) GluDTR. N=3 animals, n=11 islets for each condition.

**Fig. S5:**
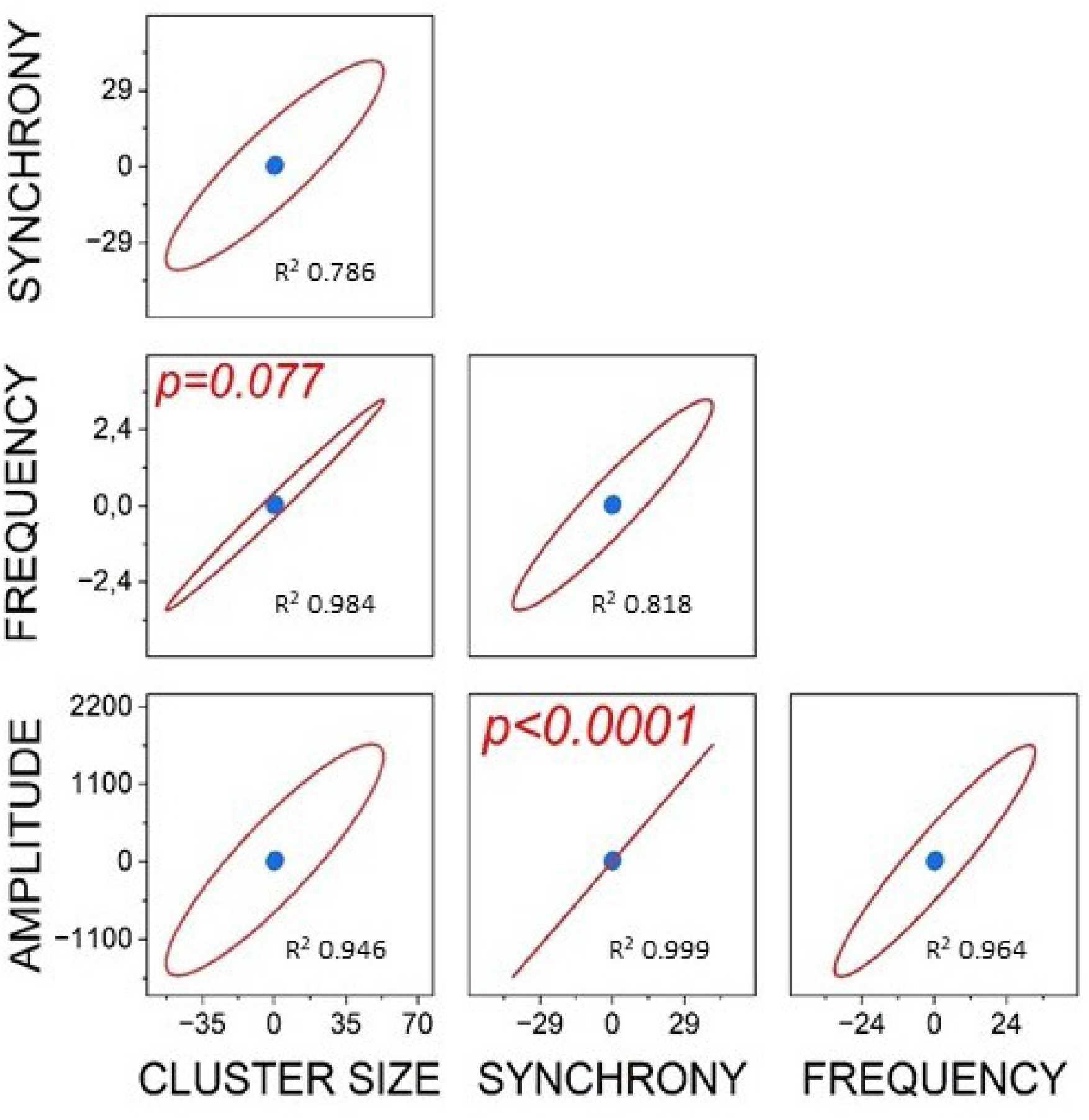
Correlations between synchrony, cluster size, frequency and amplitude. The data from HD-MEA recordings in terms of mean frequency or mean amplitude during G8.2 stimulation (in the presence of amino acids) of wild type islets were used. Pearson correlations with center points and 99% confidence intervals (red line), R^2^ and relevant p values are given; Data from 3 animals, and 11 islets total as in Fig. 5.

**Fig. S6:**
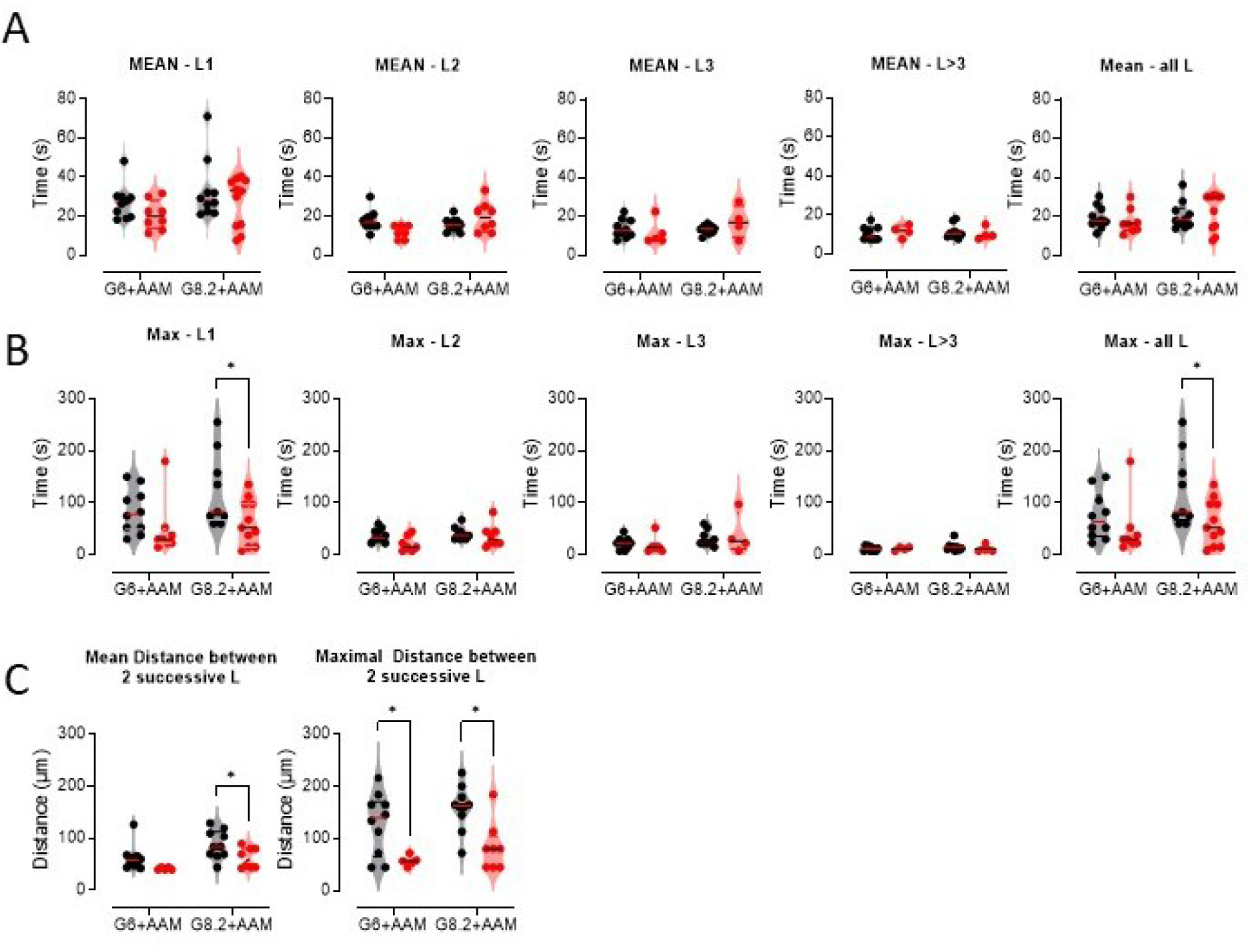
Temporal stability of leader regions and distances between successive leader regions. (**A)** mean stability in seconds for first, second, third and subsequent leader regions as well as all leader regions. Black, wild type; red, GluDTR. (**B**) Maximal stability for leader regions. (**C**) Mean and maximal distance between two successive leader regions. Original data as in Fig 6. 2-way ANOVA and Kruskal; *, 2p<0.05.

